# The role of agentive and physical forces in the neural representation of motion events

**DOI:** 10.1101/2023.07.20.549905

**Authors:** Seda Karakose-Akbiyik, Oliver Sussman, Moritz F. Wurm, Alfonso Caramazza

**Affiliations:** Department of Psychology, Harvard University, Cambridge, Massachusetts, USA; Center for Mind/Brain Sciences – CIMeC, University of Trento, Rovereto, Italy; University of Pompeu Fabra, Spain

## Abstract

How does the brain represent information about motion events in relation to agentive and physical forces? In this study, we investigated the neural activity patterns associated with observing animated actions of agents (e.g., an agent hitting a chair) in comparison to similar movements of inanimate objects that were either shaped solely by the physics of the scene (e.g., gravity causing an object to fall down a hill and hit a chair) or initiated by agents (e.g., a visible agent causing an object to hit a chair). Using fMRI-based multivariate pattern analysis, this design allowed testing where in the brain the neural activity patterns associated with motion events change as a function of, or are invariant to, agentive versus physical forces behind them. Cross-decoding revealed a shared neural representation of animate and inanimate motion events that is invariant to agentive or physical forces in regions spanning frontoparietal and posterior temporal cortices. In contrast, the right lateral occipitotemporal cortex showed higher sensitivity to agentive events, while the left dorsal premotor cortex was more sensitive to information about inanimate object events that were solely shaped by the physics of the scene.

## Introduction

Understanding others’ actions is fundamental to our everyday lives. Whether navigating through a crowded street or talking to someone, our brains process a variety of cues related to people, objects, and their interactions to arrive at a meaningful interpretation. Previous work identified certain frontoparietal and posterior temporal brain regions that are involved in understanding others’ actions (Caspers et al., 2010; Watson et al., 2013; Urgesi et al., 2014; Hardwick et al., 2018). These regions encode various aspects of human actions such as body motion (Grosbras et al., 2012; Han et al., 2013), agency (Gao et al., 2012; Scholl and Gao, 2013), sociality (Iacoboni and Dapretto, 2006; Isik et al., 2017; Wurm et al., 2017), motor representations (Rizzolatti and Craighero, 2004; Calvo-Merino et al., 2006), and goals (Iacoboni et al., 2005; Cavallo et al., 2016; Patri et al., 2020).

While it is important to investigate how the brain represents information specific to human actions, it is also crucial to acknowledge that actions can be understood at a more basic level as the movements of physical objects. After all, humans are tangible objects existing in a physical world, generating forces, and moving through space. That is, the actions of an animate being can be described with respect to agency or goals, but also at a level specifying kinematics of movement (Zacks et al., 2006; Dayan et al., 2007; Mulliken et al., 2008; McAleer et al., 2014), inter-object relations (Hafri and Firestone, 2021) or the amount of physical force exerted (Liu et al., 2017). Is there a neural representation of events that encodes such properties regardless of animacy or agency? Which brain regions are more sensitive to agentive versus physical event dynamics? These are the questions addressed in the current study.

We define motion events as changes in an object’s position relative to its surroundings. For a motion event to occur, a force needs to be applied upon an object, causing it to accelerate or decelerate. Various physical forces can shape motion, and importantly, animate agents can generate and control their own movement. Relatedly, for the purposes of this study, we categorized forces that control movement as *agentive* or *physical*. For instance, a person rising from a chair involves an agentive force, controlled by the individual, in a way that cannot be reduced to external physical forces. In contrast, an inanimate object, incapable of self-propelled motion, requires an external energy source to move. The movements of an inanimate object can be fully explained by external physical forces, the source of which can be agentive (e.g., a person throwing a rock) or physical (e.g., a rock falling off a cliff).

Within this framework, our study addressed the contributions of agentive and physical forces on the neural representation of motion events. Our experimental conditions included events driven entirely by the physics of the scene devoid of any agent involvement (e.g., a ball rolls down a slope due to gravitational pull and bounces over a chair), as well as events tied to animate agents, either as the cause of inanimate motion (e.g., a visible agent causes a ball to descend a slope and bounce over a chair) or as executors of an action (e.g., an agent bounces over a chair). We ensured that the unfolding of events, motion trajectories, and inter-object relations were analogous across all experimental conditions (e.g., X bounces over Y).

Using fMRI-based multivariate pattern analysis and cross-decoding, we identified a neural representation of motion events that is invariant to agentive or physical forces in frontoparietal and posterior temporal regions that are associated with human action recognition. Furthermore, right lateral occipitotemporal cortex showed greater sensitivity to events involving animacy and agency, while the left dorsal premotor cortex was more sensitive to information about inanimate object events that were shaped by the physics of the scene. Overall, our study provides new insights into the functional properties of brain regions that are involved in human action understanding.

## Materials and Methods

### Participants

Twenty-nine healthy adult participants completed the study (nine male; age range: 21-34; *Mean*_age_: 25.62). Neuroimaging data and participant performance were inspected for quality and four subjects were excluded from the analyses (see Preprocessing for more detail). The final dataset included 25 participants. All participants were right-handed and had normal or corrected-to-normal vision. Participants provided informed consent prior to the experiment and were paid 75 dollars for a 2-hour scanning session. The experimental protocol was approved by Harvard University’s Committee on the Use of Human Subjects.

### Stimuli

To identify where in the brain the neural activity patterns associated with motion events are invariant to, or change as a function of, agentive and physical forces, we used fMRI while participants viewed 2-sec animated videos (see Figure 1). We produced the videos using Blender v2.92, a free and open-source animation software (Blender Foundation, 2021), and presented them at the center of fixation with a frame rate of 30 frames per second. For stimulus presentation, response collection, and synchronization with the scanner, we used MATLAB Psychtoolbox-3.

**Figure 1.**
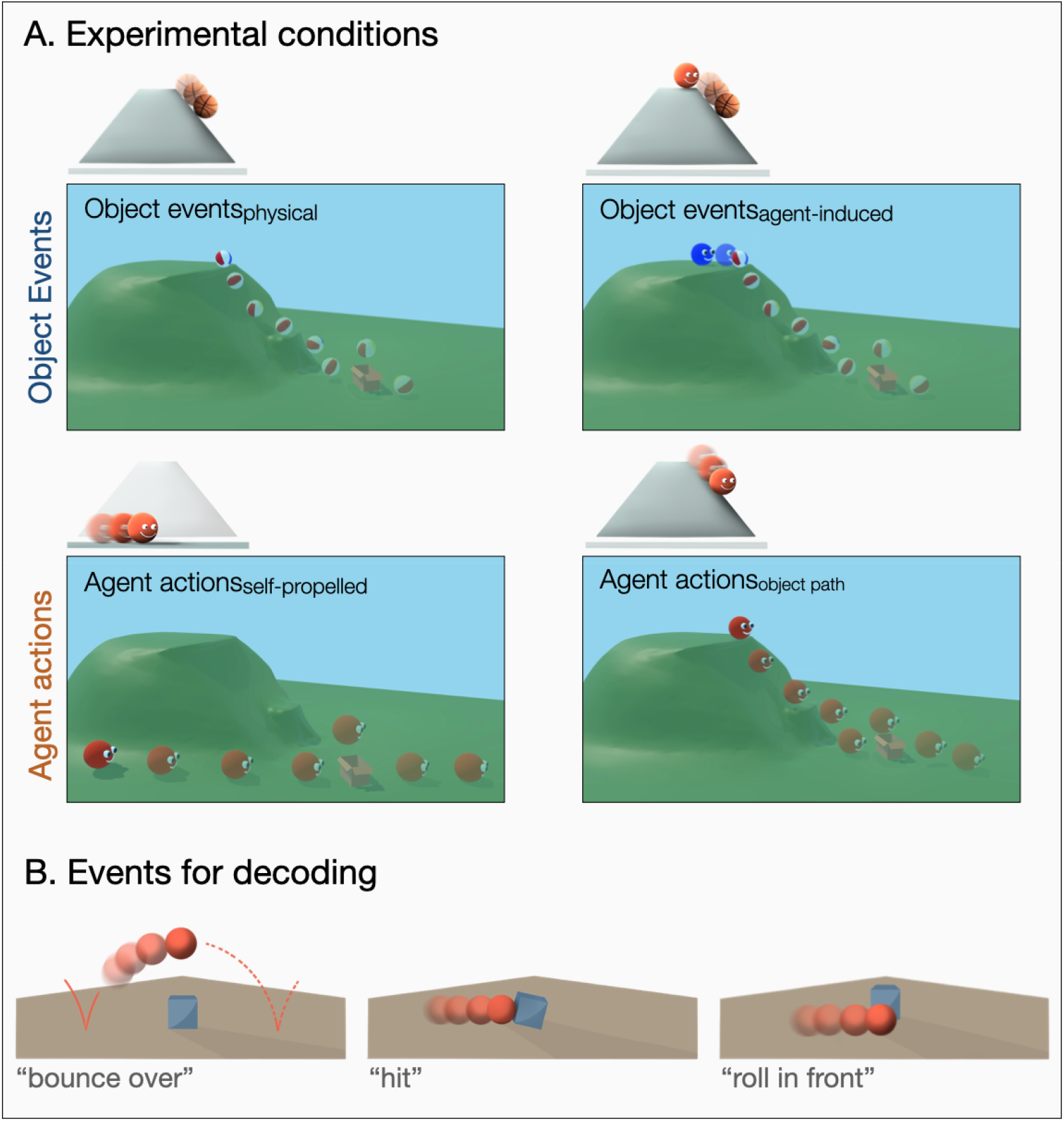
Sample stimuli and experimental design. **(A)** Experimental conditions and sample trials. For all experimental conditions, 2-sec videos were used depicting the movements of a spheric agent or a ball. **(B)** Events used for decoding. For all experimental conditions, three motion trajectories were used in relation to an animate or inanimate passive patient: bounce over – hit – roll in front.

Our stimuli comprised four event conditions: object events_physical_ (e.g., gravity makes a ball roll down a hill and the ball bounces over a chair as governed solely by the inherent physics of the scene), object events_agent-induced_ (e.g., an agent causes a ball to roll down a hill and the ball bounces over a chair), agent actions_self-propelled_ (e.g., a stationary agent at the bottom of the hill starts moving and jumps over a chair in a fully self-propelled way), and agent actions_object-path_ (e.g. an agent slides down a hill and bounces over a chair following the same trajectory as the inanimate object events, see Figure 1A for sample stimuli, see Supplementary Figure 1 for univariate activation maps). All events took place within a scene layout of a hill and a meadow. The hill in the scene enabled introducing gravity and control of physical forces such that an object can move without agent involvement (see Results for more detail on how the different conditions were used to address different aspects of motion event representation in the brain).

All experimental conditions depicted three motion trajectories with respect to a passive patient in the scene (“bounce over”, “hit”, “roll in front of”, see Figure 1B), creating 12 unique motion events (4 experimental conditions * 3 motion trajectories). These three motion trajectories were used as the basis for all decoding analyses within or across experimental conditions (for more detail, see Whole-brain searchlight MVPA). We created 64 exemplars per unique motion event, through which we introduced significant perceptual variability (see Supplementary Figure 2). Through these exemplars, all motion events were presented across four viewing angles, two moving directions (left or right), two subjects (ball-basketball for inanimate objects; red agent-blue agent for animate agents), and four passive patients (two inanimate patients: chair or box; two animate patients: pink agent-orange agent). For instance, the “roll in front of” event featuring the “object events_physical_” condition was depicted across these 64 different exemplars. This strategy helped control for low-level visual confounds that might distinguish one motion trajectory from another (e.g., presence of an animate entity, presence of occlusion, where in the scene there is movement), ensuring that decoding is not merely a consequence of such low-level visual features.

### Design of the fMRI experiment

Participants underwent one scanning session that started with an anatomical scan followed by eight functional runs; each run contained four blocks, and each block contained 28 trials (24 experimental trials, and four catch trials). We used an event-related design to present the stimuli, and the different event conditions and motion trajectories were interspersed within runs in a randomized fashion. There were 96 experimental trials (24 experimental trials per block × 4 blocks = 96) and 16 catch trials per run, and each trial consisted of a 2-sec video followed by a 1-sec fixation period. We showed longer fixation periods of 10-sec before runs, 16-sec after runs, and 10-sec between blocks. Over the course of the 96 experimental trials within a run, we showed each of the 12 unique motion events (4 event conditions * 3 motion trajectories) eight times (twice per block, once with animate and once with inanimate patients). Due to logistical challenges, some participants were not able to complete all eight runs; however, since different event conditions were balanced within runs and exemplars were sampled randomly across runs, this is unlikely to have resulted in confounds. In the final dataset, all 25 subjects provided data at least for seven out of the eight runs.

### Task

To ensure that participants paid attention to the events depicted in the videos, we conducted a catch trial detection task: participants were asked to press a button when they detected aberrant videos (16 catch trials per run, 14% of all trials). These aberrations were either perceptual, in which a visual oddity was introduced to the video (e.g., color change, freezing), or conceptual, in which the video depicted a meaningfully different movement (e.g., a ball rolling down the back of the hill; an agent turning around prior to movement). The catch trial task ensured that participants paid attention to the stimuli, both to their visual features (through the perceptual catch trials), and their higher-level aspects (through the conceptual catch trials). Responses made prior to the end of the 1-sec fixation period following each trial were counted. Prior to the scanning session, we showed participants demo videos, explained that catch trials would contain either perceptual or conceptual aberrations, and then showed a sample of the experimental layout. During the anatomical scan, participants completed a practice run, in which the screen displayed feedback after both correct and incorrect responses; no feedback was shown during the actual experiment.

During the functional scans, we presented an equal number of catch trials for each of the four experimental conditions, and the variations in viewpoint and moving direction were counterbalanced. Participants showed high performance in the catch trial task with low false alarm rates (*M* = .006, *SD* = .007) and high hit rates (*M* = .953, *SD* = .036). For two participants, catch trial performance could not be recorded due to technical issues. These participants performed with high accuracy as observed throughout the data collection, thus, we chose to keep these participants in the analysis of the neuroimaging data. One run each of three participants were excluded due to off-task behavior during the scans (e.g., sleeping).

### Data Acquisition

The neuroimaging data were collected using a 3T Siemens Prisma fMRI Scanner using a 32-channel phased-array head coil. T1-weighted structural images were obtained using a 3D MPRAGE sequence (176 sagittal slices; repetition time (TR) = 2530 msec; inversion time = 1020 msec; flip angle = 7 degrees; field of view (FoV) = 256 × 256 mm; 1×1×1 mm voxel resolution). Functional images were acquired using a T2*-weighted gradient echo-planar imaging (EPI) sequence (TR = 1500 msec; echo time (TE) = 28 msec; inter slice time = 33 msec; flip angle = 70 degrees; FoV = 200 mm × 200 mm; matrix size = 66 × 66; 3×3×3 mm voxel resolution; 45 slices with 3 mm thickness and 0 mm gap).

### Preprocessing

We preprocessed and analyzed functional and anatomical data using BrainVoyager 22.4, NeuroElf Toolboxes, CoSMoMVPA, and MATLAB 2021b (Goebel, 2012; Oosterhof et al., 2016). The first four volumes of functional runs were removed to prevent T1 saturation. Preprocessing of functional data included slice time correction, three-dimensional motion correction (trilinear interpolation, the first volume of the first run of each participant was used as reference), linear trend removal, high pass filtering (cutoff frequency of three cycles), and spatial smoothing (Gaussian kernel of 8mm FWHM for univariate analyses and 3 mm FWHM for MVPA). Functional images were registered to high-resolution anatomical images, and anatomical and functional data were normalized to Talairach space. We inspected all anatomical and functional scans for data quality and excluded scans that had a maximum absolute motion greater than 3mm and a signal-to-noise ratio lower than 130. Based on these criteria, two out of the 29 participants were excluded due to low data quality as many of their runs did not meet these quality criteria, leaving limited data for analyses. Another two participants were excluded from the analyses due to logistical issues during their scan and off-task behavior. Out of the remaining 25 participants, only one run of one participant was excluded for not meeting the quality criteria for functional scans. Thus, the analyses presented in the paper contain high quality functional data with low motion and high signal-to-noise ratio.

### fMRI Data Analysis

For each participant and run, we computed a general linear model using design matrices containing 24 event predictors (separate predictors were fit for animate-inanimate patients per 12 unique motion events), plus one predictor for catch trials. Regressors were defined as boxcar functions convolved with a canonical double-gamma hemodynamic response function. Trials were modeled as epochs lasting from video onset to offset (2-sec) and the resulting reference time courses were used to fit the signal time courses of each voxel. In total, this procedure resulted in 16 beta maps per 12 unique motion event per subject.

### Whole-brain searchlight MVPA

To investigate the neural representation of motion events in relation to agentive and physical forces, we used multivariate pattern analysis techniques (MVPA). For all MVPA analyses, decoding was completed over three motion trajectories defined with respect to a passive patient (i.e., bounce over – hit – roll in front, see Figure 1B). A linear discriminant analysis classifier algorithm was trained and tested on the beta maps, divided into four-voxel-radius spheres (12mm), to classify the associated stimuli by motion trajectory (three-way decoding, chance-level: 33.33%).

To investigate the neural representation of motion events in relation to different experimental conditions, we employed two decoding approaches. Firstly, within-condition decoding was used, where a classifier was trained and tested to differentiate between the three motion events within a specific experimental condition (e.g., self-propelled agent actions) using leave-one-out cross-validation. To test for differences in decoding strength across the different experimental conditions, we compared the respective decoding maps using whole-brain two-tailed paired t-tests. To identify the neural representations shared across agentive and physical dynamics, we performed cross-decoding. This involved training the classifier on data from one condition (e.g., distinguishing bounce over – hit – roll in front for physical object events) and then testing it on data from another condition (e.g., distinguishing bounce over – hit – roll in front for self-propelled agent actions). We repeated this process in the opposite direction and averaged the resulting classification accuracies. The resulting accuracy maps from the decoding analyses were entered into one-tailed t-tests to test for above chance classification in the whole brain (chance level: 33.33%). We used the Monte Carlo Cluster based method to correct for multiple comparisons (initial threshold: *p* = .001, 10000 simulations), and visualized the results on a cortex-based surface.

In our decoding analyses, we treated the motion events with different subjects, animate or inanimate passive patients, viewing angles, and moving directions as the “same event”. This approach allowed the classifier to learn to detect the motion events regardless of variability in these factors. Since the classifiers were trained to distinguish the three motion events across these variations, any differences observed in decoding between different experimental conditions (e.g., between object events_physical_ versus agent actions_self-propelled_) cannot be reduced to these factors.

### ROI analysis

We primarily focused on and report whole-brain searchlight maps. However, to gain a more detailed understanding of how information about motion events is represented in different brain regions, we conducted region-of-interest (ROI) analyses on specific areas that are traditionally associated with human action observation following a meta-analysis (Caspers et al., 2010): the lateral occipitotemporal cortex (LOTC), inferior parietal lobule (IPL), dorsal premotor cortex (PMd), ventral premotor cortex (PMv), posterior superior temporal sulcus (pSTS), and superior parietal lobule (SPL). The MNI coordinates from the meta-analysis were converted to TAL coordinates using Yale Bioimage Suite (Papademetris et al., 2006; Lacadie et al., 2008). Since the meta-analysis provided a different number of ROIs in the frontoparietal cortices for left and right hemispheres, for simplicity, we used the centroid of Brodmann areas 6 and 7 for dorsal premotor cortex and superior parietal lobules, respectively (Lacadie et al., 2008). All ROIs were created as spheres with a 12mm radius around their respective coordinates (TAL coordinates: left LOTC [-45 -71 6], left IPL [-58 -23 34], left PMd [-28 0 48], left PMv [-48 8 29], left pSTS [−52 −49 11], left SPL [-18 -57 50]; right LOTC [52 -63 5], right IPL [44 -31 41], right PMd [28 1 47], right PMv [50 12 27], right pSTS [54 −40 8], right SPL [24 -56 54]).

For the ROI analyses, we extracted decoding accuracies from the searchlight maps for each classification scheme (e.g., decoding of self-propelled agent actions), participant, and ROI. We then entered the decoding accuracies for ROIs into FDR-corrected one-tailed t-tests to identify which ROIs showed above chance classification. To investigate differences in decoding strength across event types and ROIs, we applied linear mixed effects models using the *lme4* package (Bates et al., 2015) in R version 4.1.1. (R Core Team, 2021). To examine interactions between event type and ROI, we compared models with and without the interaction term using a likelihood ratio test. For instance, to test the interaction between event type and ROI, Model 1 included Classification Accuracy, Region, Event Type, and Subject ID as a random effect (1 | Subject ID) and did not allow for an interaction term between Region and Event Type (Region + Event Type). We then compared Model 1 with Model 2, which expanded on Model 1 by including an interaction term between Region and Event Type (Region * Event Type). After significant interactions or main effects, we conducted FDR corrected post-hoc two-tailed tests of estimated marginal means to investigate which conditions are responsible for driving the observed effects.

## Results

### A shared neural representation of motion events across agentive and physical forces

One of our main aims was to identify a neural representation of motion events that is invariant to agentive or physical forces behind them. To this end, we first focused on two conditions: self-propelled agent actions and physical object events. To make sure that the self-propelled agent actions depicted full agentive control, an animated agent (a solid-color sphere with eyes and mouth) stood stationary on a flat surface and then started moving (see Figure 1A). These agent actions served as reference for human actions that are broadly studied in the literature, as substantial literature has shown that animations of simple geometric figures can elicit robust perceptions of animacy and agency, especially when they move in a self-propelled way (Heider and Simmel, 1944; Michotte, 1946; Scholl and Tremoulet, 2000; Blakemore et al., 2001).

On the opposite end of these self-propelled, agentive actions were physical object events that were governed fully by the physics of the scene without any agent involvement. These events started with a ball toppling off a ledge atop a hill. The ball would then fall down the hill to complete one of the three motion trajectories (i.e., bounce over – hit – roll in front) as controlled by a physics engine built into the animation software. An independently collected behavioral survey confirmed that observers did not attribute agent involvement to these events (see Supplementary Figure 3, Supplementary Note 1).

To identify brain regions that encode a general neural representation of motion events independent of agentive or physical forces, we conducted cross-decoding MVPA across self-propelled agent actions and physical object events. Cross-decoding revealed robust generalization across self-propelled agent actions and physical object events in various frontoparietal and posterior temporal clusters in both hemispheres (see Figure 2A, for ROI analysis see Supplementary Figure 4B). Success in this cross-decoding reflects event representations that are not tied to animacy or the type of force controlling the event – agentive or physical.

**Figure 2.**
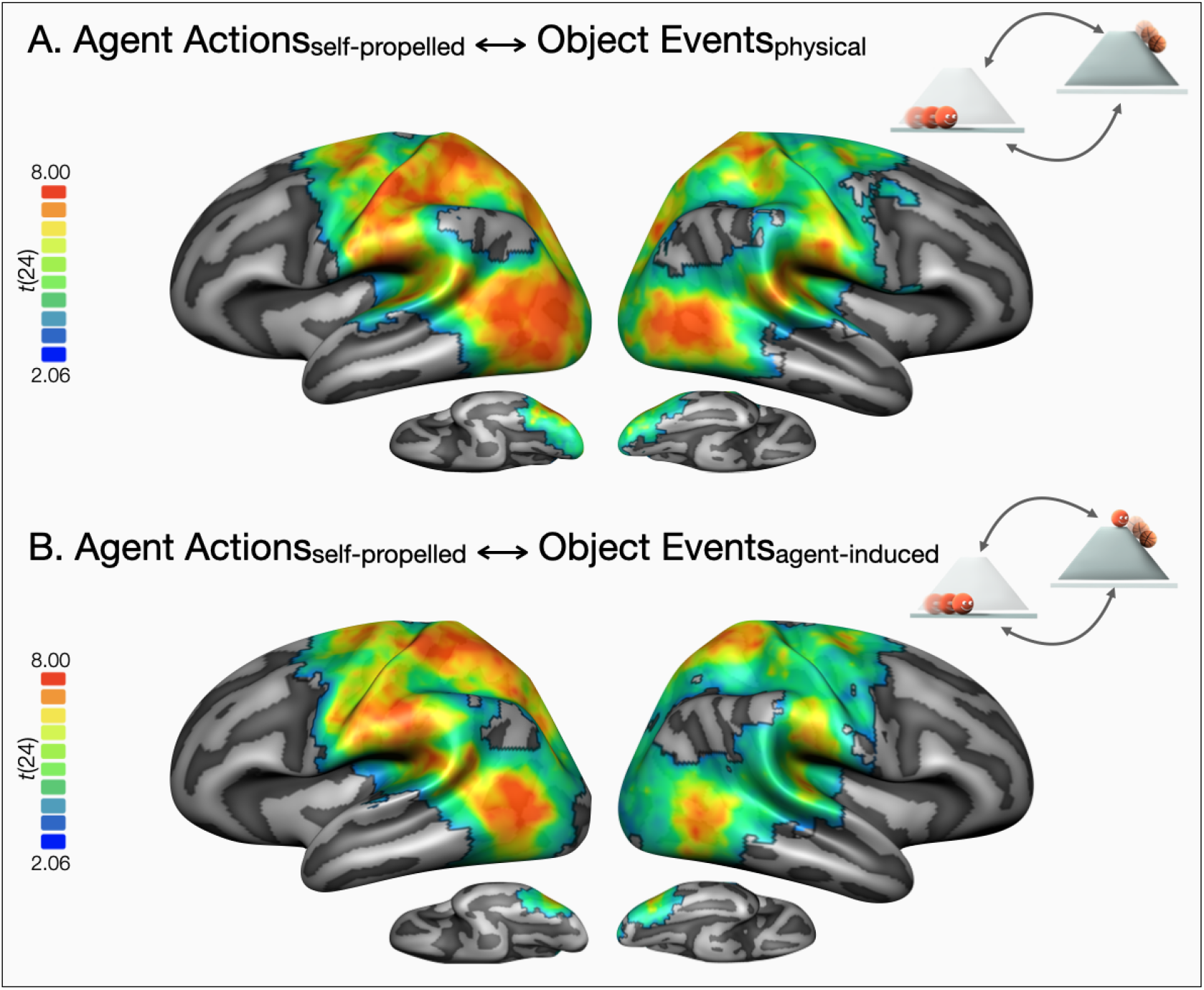
Decoding of observed events by generalizing across animacy, and agentive and physical forces. **(A)** Results of whole-brain three-way decoding searchlight across self-propelled agent actions and physical object events. **(B)** Results of whole-brain three-way decoding searchlight across self-propelled agent actions and agent-induced object events (one tailed t-tests against chance level 33.33%). Correction for multiple comparisons was conducted using the Monte Carlo Cluster based method (*p*_initial_ = .001). Only the areas that survived correction are presented as highlighted by a black border.

We found a shared neural representation for self-propelled agent actions and physical object events in regions classically associated with human action recognition. As we noted before, the source of an inanimate objects’ movement can be agentive or physical. Is this shared neural representation we identified independent of whether the movement is caused by an agent or physical forces? How does the presence of causal agency in the movement of inanimate objects affect their neural representation compared to actions performed by animate entities?

To address this question, we created instrumental object events that had a visible agent cause behind them: agent-induced object events. Here, a visible agent pushed a ball down the hill and as a result, the ball bounced over – hit – rolled in front of a passive patient (see Figures 1A-B). The movements of the initiator agent were always the same across the three motion trajectories, and the meaningful distinctions happened in relation to the resulting movements of the ball. Thus, successful decoding of the three motion trajectories could not rely on the initiator agent’s movement and should capture the movements of the ball. Furthermore, to control for possible perceptual confounds between the physical object events and agent-induced object events, the ball’s movements were made identical in the two conditions: all that differed was the presence of an animate agent initiating the object motion in agent-induced object events, and the rest of the path was controlled by the physics engine. An independently collected behavioral survey showed that observers endorsed the involvement of agents in these agent-induced object events (see Supplementary Figure 3, Supplementary Note 1).

Cross-decoding of self-propelled agent actions and agent-induced object events was successful in overlapping frontoparietal and posterior temporal clusters (see Figure 2B). We hypothesized that regions that are sensitive to causal agency might demonstrate greater generalization between agent actions and agent-induced object events compared to physical object events that were purely governed by physical forces. No reliable differences were found in cross-decoding strength across the whole-brain when comparing the two cross-decoding maps from self-propelled agent actions to agent-induced or physical object events. (see Supplementary Figure 4A). In the ROI-analysis, cross-decoding between self-propelled agent actions and physical object events was stronger than that between self-propelled agent actions and agent-induced object events in right and left LOTC (see Supplementary Figure 4B, Supplementary Note 2). However, note that agent-induced object events had a more complex event structure and depicted instrumental actions, whereas physical object events and self-propelled agent actions did not. This difference in complexity might explain better cross-decoding of self-propelled agent actions to physical object events compared to that between self-propelled agent actions and agent-induced object events in bilateral LOTC and might have masked any possible contributions of causal agency to cross-animacy generalization.

### Searching for signs of agency and animacy in the neural representation of motion events

Cross-decoding showed that regions classically associated with human action recognition carry a shared neural code for motion events that generalizes across animate and inanimate entities, and agentive and physical forces. However, agent actions and movements of inanimate objects are marked by key differences. For example, their kinematics may differ, with agents able to change direction and speed while inanimate object movement is determined by external physical forces. Additionally, agent actions reflect the goal-directed behavior of animate entities, while object events pertain to inanimate physical objects. There is evidence that sensitivity to animacy and agency is present even from infancy (Opfer, 2002; Csibra, 2003, 2008; Tremoulet and Feldman, 2006; Liu et al., 2017) and animacy is a widely established principle of organization in the neural representation of objects (Konkle and Caramazza, 2013; Grill-Spector and Weiner, 2014; Peelen and Downing, 2017; Wurm and Caramazza, 2022). Given these considerations, we next asked if any brain regions are sensitive to agentive versus physical forces while encoding dynamic event information. To address the differential roles of agentive and physical forces on the neural representation of motion events, we completed different decoding analyses. Specifically, we trained and tested classifiers with neural activity patterns associated with different experimental conditions separately (i.e., within self-propelled agent actions or physical object events), and compared their decoding strengths.

We first investigated the decoding strengths of self-propelled agent actions and physical object events, which provides a robust testbed to investigate the relative contributions of agentive and physical forces to the neural representation of motion events. In line with previous findings, both event types were decoded in regions spanning posterior temporal, frontal, and parietal cortices (see Figures 3A-B). Comparing the two event types, a two-tailed whole-brain paired t-test revealed multiple clusters spanning the right lateral occipitotemporal cortex, posterior middle temporal sulcus, temporoparietal junction, and supramarginal gyrus that can better distinguish self-propelled agent actions compared to physical object events (see Figure 3C). Although additional clusters in left posterior temporal cortex showed better decoding of self-propelled agent actions, and some clusters in left dorsal premotor cortex showed better decoding of physical object events (*p*s < .005), these effects did not survive correction for multiple comparisons in the whole brain. For a more fine-grained analysis, we turned to the ROI analysis (see Figure 3D).

**Figure 3.**
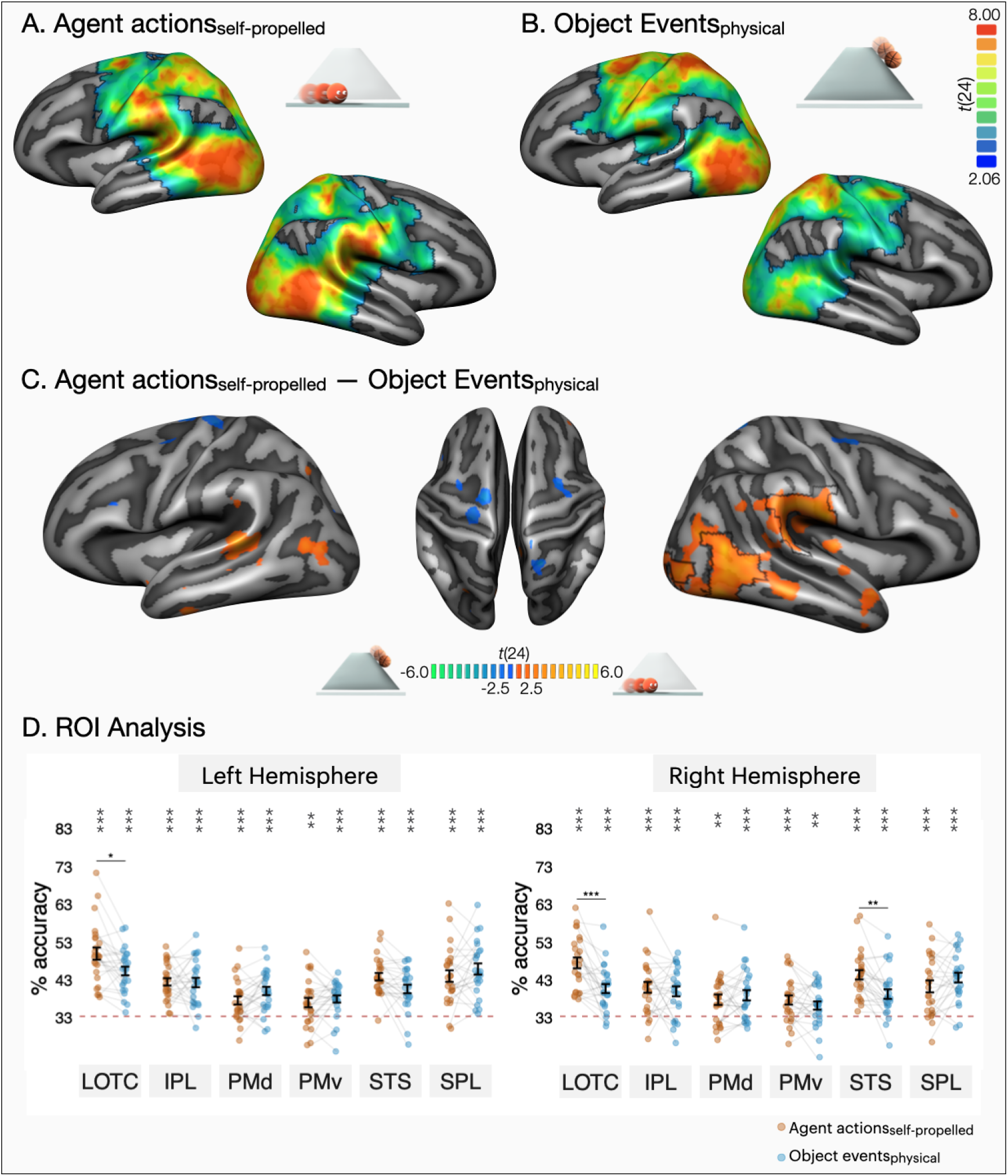
Decoding of self-propelled agent actions and physical object events. Results of whole-brain three-way decoding searchlight for (**A)** self-propelled agent actions and **(B)** physical object events (one tailed t-tests against chance 33.33%). Correction for multiple comparisons was conducted using the Monte Carlo Cluster based method (*p*_initial_ = .001). Only the areas that survived correction are presented as highlighted by a black border. (**C)** Two-tailed whole brain decoding contrast of self-propelled agent actions physical object events. Black outlines mark areas that survived Monte Carlo Cluster based correction (*p*_initial_ = .001). The map is thresholded at *p* < .02 to demonstrate significant differences that do not survive correction for multiple comparisons. (**D)** ROI decoding accuracies for self-propelled agent actions and physical object events. Error bars indicate standard error of the mean (SEM), and asterisks indicate FDR-corrected effects of one-tailed t-tests for comparisons against chance level (33.33%, \**p* < .05, \*\**p* < 0.01, \*\*\**p* < 0.001). Individual participants are connected via light gray lines. FDR-corrected pairwise two-tailed tests of estimated marginal means showed better decoding of self-propelled agent actions in left and right LOTC, and right pSTS (\**p* < .05, \*\**p* < 0.01, \*\*\**p* < 0.001).

To compare decoding accuracies of self-propelled agent actions and physical object events across different ROIs within each hemisphere, we fitted linear mixed effect models testing the interaction of event type and ROI. This ROI by event type interaction was significant both in the left hemisphere (χ2[5] = 32.23, *p* < .001, ΔAIC = 22.23) and in the right hemisphere (χ2[5] = 42.30, *p* < .001, ΔAIC = 32.30). Post-hoc contrasts revealed that self-propelled agent actions were decoded at a higher accuracy than physical object events in left LOTC (*b* = 4.60, *p* = .020, *d* = .68). In the remainder of the left hemisphere ROIs, self-propelled agent actions and physical object events were classified with a comparable accuracy (IPL: *b =*.19, *p* = .905, *d* = .03; PMd: *b* = -2.49, *p* = .217, *d* = -.44; PMv: *b* = -.95, *p* = .646, *d* = -.16; pSTS: *b* = 3.25, *p* = .111, *d* = .59; SPL: *b* = -1.87*, p* = .342, *d* = -.23). In the right hemisphere, self-propelled agent actions were decoded at a higher accuracy than physical object events in right LOTC (*b* = 6.79, *p* < .001, *d* = .94) and pSTS (*b* = 5.10, *p* = .002, *d* = .81). Self-propelled agent actions and physical object events were classified with comparable accuracy in the remainder of the right hemisphere ROIs (IPL: *b* = 1.06, *p* = .476, *d* = .13; PMd: *b* = -1.17, *p* = .476, *d* = -.17; PMv: *b* = 1.48, *p* = .476, *d* = .23; SPL: *b* = -2.33, *p* = .234, *d* = -.32).

Stronger decoding of self-propelled agent actions in LOTC and pSTS, particularly in the right hemisphere, replicates our previous study that addressed the shared and distinct neural representations of observed human actions and inanimate object events (Karakose-Akbiyik et al., 2023). The current study adds to these findings and highlights that explicit body motion, and the associated movement kinematics, are not necessary for driving differences between animate and inanimate movement in these regions.

### Comparing agent actions and object events in a more controlled setting

Self-propelled agent actions presented so far were highly effective in giving impressions of intentionality and agentive control. However, the overall trajectories of these stimuli were not completely matched with that of physical object events. In physical object events, the ball first fell down a hill, and completed one of the three motion trajectories. The agent actions on the other hand, started on a meadow and did not complete the extra step of going down a hill (see Figure 1A). To compare agent actions and object events in a more controlled setting, we created another agent action condition: agent actions_object-path_. These stimuli started with an agent standing atop a hill. The agent then slid down the hill to complete one of the three motion trajectories. Thus, the motion trajectories of the agent actions_object-path_ stimuli were matched with that of physical object events, providing a controlled setting to identify the contributions of animacy. Furthermore, like physical object events, the motion trajectories of agent actions_object-path_ were also determined by the physics engine. An independently collected behavioral survey ensured that these agent actions were still perceived as agentive compared to physical object events validating the use of these stimuli to test the contributions of agentive and physical forces to the neural representation of event dynamics (see Supplementary Figure 3, Supplementary Note 1).

Consistent with previous results, decoding of agent actions_object-path_ and object events was successful in mostly overlapping frontoparietal and posterior temporal brain regions (see Figure 4A-B). Comparing the decoding strengths of the two conditions in the whole brain, a two-tailed t-test revealed better decoding for physical object events in a left dorsal premotor cortex cluster (see Figure 4C). Additional clusters in bilateral SPL and right LOTC showed a significant difference between the two event types (*p*s < .005), but these effects did not survive correction for multiple comparisons in the whole brain. For a more fine-grained analysis, we again turned to ROIs (see Figure 4D).

**Figure 4.**
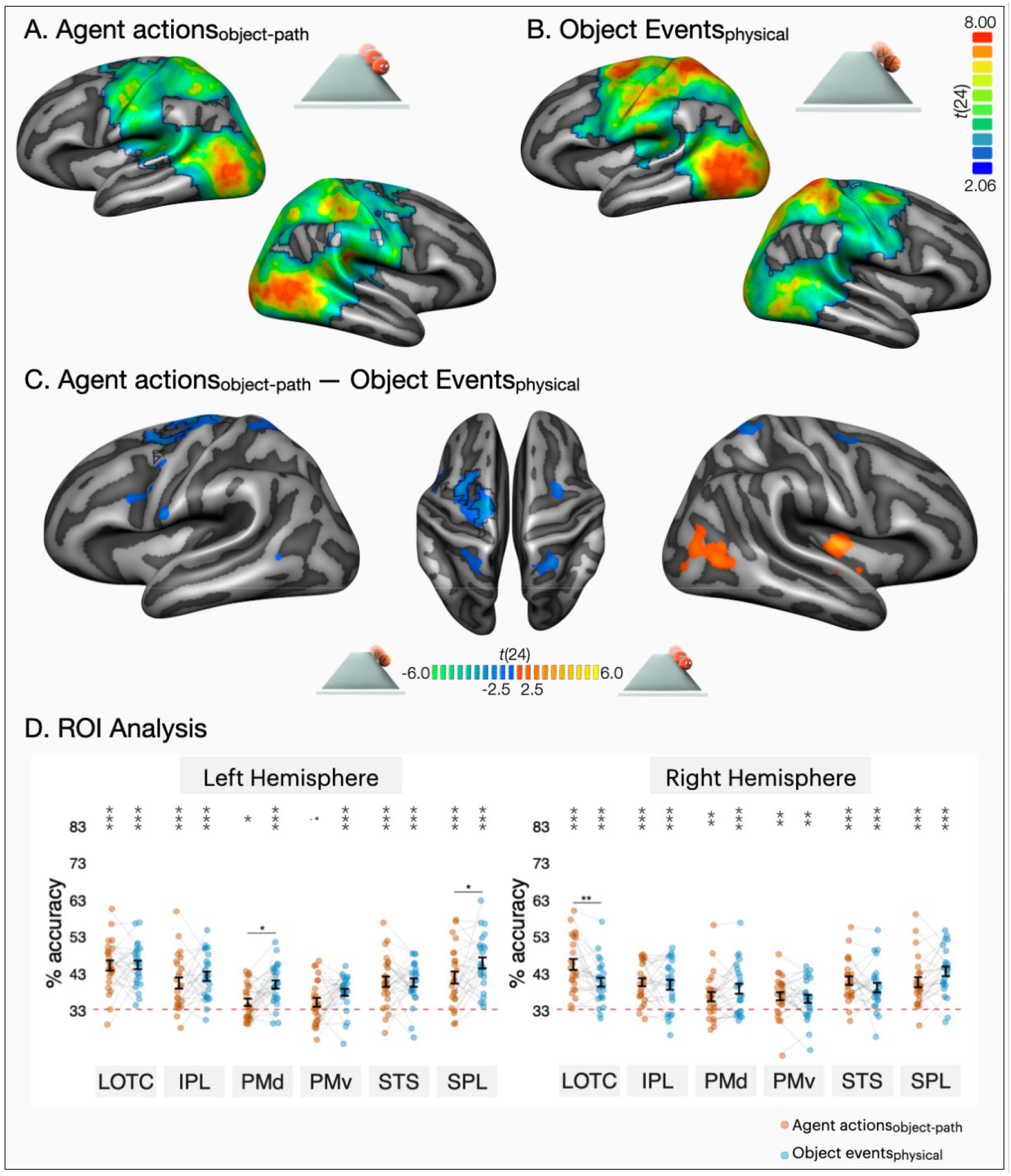
Decoding of agent actions_object-path_ and physical object events. Results of whole-brain three-way decoding searchlight for (**A)** agent actions_object-path_ and **(B)** physical object events (one tailed t-tests against chance 33.33). Correction for multiple comparisons was conducted using the Monte Carlo Cluster based method (*p*_initial_ = .001), and only the regions that survived correction for multiple comparisons are presented as highlighted by a black border. (**C)** Two-tailed whole brain t-test contrast of agent actions_object-path_ and physical object events decoding. Black outlines mark clusters that survived Monte Carlo Cluster based correction (*p*_initial_ = .001). The map is thresholded at *p* < .02 to demonstrate significant differences that do not survive correction. (**D)** ROI decoding accuracies for agent actions_object-path_ and physical object events. Error bars indicate standard error of the mean (SEM), and asterisks indicate FDR-corrected effects of one-tailed t-tests for comparisons against chance level (33.33%, \**p* < .05, \*\**p* < 0.01, \*\*\**p* < 0.001). Individual participants are connected via light gray lines. FDR-corrected pairwise two-tailed tests of estimated marginal means showed better decoding of agent actions_object-path_ in right LOTC, and better decoding of physical object events in left PMd and SPL (\**p* < .05, \*\**p* < 0.01, \*\*\**p* < 0.001).

All ROIs showed above chance decoding of agent actions_object-path_ except for left ventral premotor cortex, which bordered at significance (*p* = .06). To test which regions show a difference in decoding for the two event types, we fitted linear mixed effect models testing the interaction of event type and ROI. This ROI by event type interaction was significant both in left (χ2[5] = 23.59, *p* < .001, ΔAIC = 13.59) and right hemispheres (χ2[5] = 34.40, *p* < .001, ΔAIC = 24.40). In the left hemisphere, physical object events were decoded at a higher accuracy than agent actions_object-path_ in left dorsal premotor cortex (*b* = -4.79, *p* = .017, *d* = -.65) and superior parietal lobule (*b* = -3.92, *p* = .043, *d* = -.43). Agent actions_object path_ and physical object events were classified with comparable accuracy in the rest of the left hemisphere ROIs (LOTC: *b* = - .15, *p* = .923, *d* = -.02; IPL: *b* = -1.83, *p* = .378, *d* = -.21; PMv: *b* = -2.69, *p* = .184, *d* = -.36; pSTS: *b* = .24, *p* = .923, *d* =.03). In the right hemisphere, agent actions_object-path_ were decoded at a higher accuracy than physical object events in LOTC (*b* = 4.78, *p* = .003, *d* = .65). The rest of the right hemisphere ROIs showed comparable decoding of agent actions_object-path_ and physical object events (IPL: *b* = .71, *p* = .607, *d* = .10; PMd: *b* = -2.13, *p* = .243, *d* = -.35; PMv: *b* = .71, *p* = .607, *d* = .13; pSTS: *b* = 1.85, *p* = .267, *d* = .26), with a marginal trend in right SPL (*b* = -2.94, *p* = .099, *d* = -.45).

Overall, we found increased sensitivity to events signaling animacy and agency in right lateral occipitotemporal cortex and posterior superior temporal sulcus. We also found greater sensitivity to physical event dynamics in left dorsal premotor cortex and, in the ROI analyses, also superior parietal lobule. Increased sensitivity to physical as opposed to agentive event dynamics in superior parietal lobule also held when comparing physical and agent-induced object events (see Supplementary Figure 5, Supplementary Note 3). Overall, compared to events that had some agent involvement, motion events that followed the physics of the scene emphasized parts of dorsal premotor cortex and superior parietal lobule.

## Discussion

In frontoparietal and posterior temporal brain regions associated with human action understanding, we identified a shared neural representation of animate and inanimate motion that is also invariant to agentive or physical forces shaping the event dynamics. The right LOTC and pSTS exhibited higher sensitivity to events signaling animacy and agency, while the left dorsal premotor cortex and superior parietal lobules were more sensitive to events that were shaped by the inherent physics of the scene. Together with recent work showing a shared neural code for animate and inanimate motion, our findings highlight the general role of frontoparietal and posterior temporal regions in encoding the physics and kinematics of events regardless of animacy (Albertini et al., 2021; Karakose-Akbiyik et al., 2023). Here, we directly assessed the contributions of agentive versus physical control over movement and showed that these regions can encode information about motion events at a level that does not concern whether the source of movement is tied to an agentive or physical force.

We found increased sensitivity to events with agent involvement in right LOTC and pSTS. This finding is in line with the previous literature. Overall, it has been widely established that right LOTC and pSTS are sensitive to certain human-specific aspects of actions such as biological motion, social interactions, agency and intentionality (Saxe et al., 2004; Grafton and Hamilton, 2007; Isik et al., 2017; Tarhan and Konkle, 2020; Lee Masson and Isik, 2021; Pitcher and Ungerleider, 2021; Schultz and Frith, 2022). Nearby regions are also known to be involved in higher level cognitive processes such as theory of mind and social cognition (Pelphrey et al., 2004; Deen et al., 2015). Combined with previous findings, our results underscore that the presence of simple cues signaling animacy (e.g., facial features) and agency (e.g., self-propelled motion) are enough to drive agent-specific responses in the right LOTC and pSTS, in the absence of bodies, or when movement patterns are matched between agents and inanimate objects in terms of their interpretability.

We found increased sensitivity to events controlled by the physics of the scene in dorsal premotor cortex and superior parietal lobule. These regions are implicated in human action observation but are also considered to be a part of the brain’s *intuitive physics network* (Fischer et al., 2016; Pramod et al., 2022). There is now growing evidence that dorsal premotor cortex and superior parietal lobule are recruited when observers predict the unfolding of physical events as opposed to making other non-physical judgements about a scene (e.g., judging where an unstable tower of blocks would fall as opposed to judging whether it has more yellow or blue blocks). In these regions, there is also evidence for explicit representation of physical features such as object mass in both action planning (Van Nuenen et al., 2012) and physical inference (Schwettmann et al., 2019). The dorsal premotor cortex and superior parietal lobules have also been shown to respond more to dynamic scenes depicting higher physical content, such as moving objects, compared to faces, scenes, or moving bodies (Fischer et al., 2016). Given these previous findings, increased sensitivity to physical event dynamics in dorsal premotor cortex and superior parietal lobule might have to do with greater recruitment of parameters related to physical inference.

At this juncture, we would like to note that in a recent study, we found higher sensitivity to human actions compared to inanimate object events in the same superior parietal lobule ROI (Karakose-Akbiyik et al., 2023). However, in the current study, superior parietal lobule tended to show higher sensitivity to physical object events compared to cases where there was some animate agent involvement (e.g., agent actions, agent-induced object events). At a first glance, the reversal of the effect in SPL might be surprising. However, it is worth noting that in Karakose-Akbiyik et al. (2023), human actions depicted human body motion that has more complex mechanics than similar movements of a ball. In the current study, on the other hand, both animate agents and inanimate objects were presented as spherical geometric shapes. Furthermore, physical object events were fully governed by the physics engine and did not give any impressions of agent involvement, even behind the scenes. Taken together, these findings imply that SPL is sensitive to physical properties of movement, rather than agent-specific aspects of events. Hence, it might show sensitivity to the animate or the inanimate domain depending on the context and the specific kinds of stimuli. We do not have any direct evidence speaking to this hypothesis, but only a comparison of the patterns in our two studies. Future work could build upon these findings by investigating how the recruitment of regions associated with physical inference varies across animate and inanimate movement by using stimuli with varying physical complexity.

Overall, our findings contribute to an emerging framework on how the brain represents information about dynamic scenes. It appears that a specialized right lateralized system centered around LOTC and pSTS is particularly involved in processing information relevant for social aspects of dynamic scenes such as animacy, agency, and social interactions (Isik et al., 2017; Sliwa and Freiwald, 2017; Pitcher and Ungerleider, 2021; Dima et al., 2022). Conversely, a domain-general network comprising premotor cortex and superior parietal lobule are involved in physical inferences and prediction (Fischer et al., 2016; Yildirim et al., 2019; Fischer and Mahon, 2021).

This framework raises some interesting questions. For instance, our study revealed a shared neural code spanning various frontoparietal and posterior temporal brain regions that represents information about motion events more broadly, including those regions that showed increased sensitivity to animacy and agency (e.g., right LOTC, pSTS). What shared aspects of actions and physical object events are encoded by this overarching neural representation? While shared kinematics and spatiotemporal dynamics may be involved, further specification is needed. Moreover, both the frontoparietal and posterior temporal brain regions participate in processing of dynamic scene information. What are their distinct roles? What specific elements of dynamic scenes do they each encode? Additionally, the notion of a shared neural system that is applicable to both animate and inanimate motion presents a plausible hypothesis. However, making claims of such generality is challenging due to the limitations of fMRI measures, particularly in terms of group averaging and comparisons across different studies and paradigms. Therefore, the question persists as to whether this proposed domain-general network encompasses subcomponents that differentiate between animate and inanimate movement, paralleling distinctions observed in the object domain (Grill-Spector and Weiner, 2014; Wurm and Caramazza, 2022).

To sum up, we found a shared neural representation of animate and inanimate motion that is also invariant to agentive or physical forces in various frontoparietal and posterior temporal brain regions. Furthermore, the right lateral occipitotemporal cortex and posterior superior temporal sulcus showed higher sensitivity to cues related to animacy and agency, while the left dorsal premotor cortex and superior parietal lobules showed higher sensitivity to events that were controlled by the physics of the scene. Overall, our findings provide new insights into the contributions of agentive and physical forces in the neural representation of event dynamics and highlight the importance of a unified approach that takes both factors into account.

## Acknowledgements

We thank the members of the Cognitive Neuropsychology Lab, the Cambridge Writing Workshop, Tomer Ulman, and Talia Konkle for valuable discussions. We extend our thanks to Sena Kocyigit for her help in generating the iconography in Figure 1. Funding for this study was provided by the Center for Brain Science at Harvard University (to O.S.), Psychology Department at Harvard University (to S.K-A.), and Texas Woman’s University Woodcock Institute Research Grant (to S.K-A.).

## Data availability

Neuroimaging data, sample stimuli, and Supplementary Information are deposited at the Open Science Framework (https://osf.io/z6rfm/).

## References

Albertini D, Lanzilotto M, Maranesi M, Bonini L (2021) Largely shared neural codes for biological and nonbiological observed movements but not for executed actions in monkey premotor areas. J Neurophysiol 126:906–912.

Bates D, Mächler M, Bolker B WS (2015) Fitting Linear Mixed-Effects Models Using lme4. Journal of Statistical Software 67:1–48.

Blakemore SJ, Fonlupt P, Pachot-Clouard M, Darmon C, Boyer P, Meltzoff AN, Segebarth C, Decety J (2001) How the brain perceives causality: an event-related fMRI study. Neuroreport 12:3741–3746.

Calvo-Merino B, Grèzes J, Glaser DE, Passingham RE, Haggard P (2006) Seeing or doing? Influence of visual and motor familiarity in action observation. Curr Biol 16:1905–1910.

Caspers S, Zilles K, Laird AR, Eickhoff SB (2010) ALE meta-analysis of action observation and imitation in the human brain. Neuroimage 50:1148–1167.

Cavallo A, Koul A, Ansuini C, Capozzi F, Becchio C (2016) Decoding intentions from movement kinematics. Sci Rep 6:37036.

Csibra G (2003) Teleological and referential understanding of action in infancy. Philos Trans R Soc Lond B Biol Sci 358:447–458.

Csibra G (2008) Goal attribution to inanimate agents by 6.5-month-old infants. Cognition 107:705–717.

Dayan E, Casile A, Levit-Binnun N, Giese MA, Hendler T, Flash T (2007) Neural representations of kinematic laws of motion: evidence for action-perception coupling. Proc Natl Acad Sci U S A 104:20582–20587.

Deen B, Koldewyn K, Kanwisher N, Saxe R (2015) Functional organization of social perception and cognition in the superior temporal sulcus. Cereb Cortex 25:4596–4609.

Dima DC, Tomita TM, Honey CJ, Isik L (2022) Social-affective features drive human representations of observed actions. Elife 11 Available at: http://dx.doi.org/10.7554/eLife.75027.

Fischer J, Mahon BZ (2021) What tool representation, intuitive physics, and action have in common: The brain’s first-person physics engine. Cogn Neuropsychol 38:455–467.

Fischer J, Mikhael JG, Tenenbaum JB, Kanwisher N (2016) Functional neuroanatomy of intuitive physical inference. Proceedings of the National Academy of Sciences 113:E5072–E5081 Available at: http://dx.doi.org/10.1073/pnas.1610344113.

Gao T, Scholl BJ, McCarthy G (2012) Dissociating the detection of intentionality from animacy in the right posterior superior temporal sulcus. J Neurosci 32:14276–14280.

Goebel R (2012) BrainVoyager--past, present, future. Neuroimage 62:748–756.

Grafton ST, Hamilton AF de C (2007) Evidence for a distributed hierarchy of action representation in the brain. Hum Mov Sci 26:590–616.

Grill-Spector K, Weiner KS (2014) The functional architecture of the ventral temporal cortex and its role in categorization. Nat Rev Neurosci 15:536–548.

Grosbras M-H, Beaton S, Eickhoff SB (2012) Brain regions involved in human movement perception: a quantitative voxel-based meta-analysis. Hum Brain Mapp 33:431–454.

Hafri A, Firestone C (2021) The Perception of Relations. Trends Cogn Sci 25:475–492.

Han Z, Bi Y, Chen J, Chen Q, He Y, Caramazza A (2013) Distinct Regions of Right Temporal Cortex Are Associated with Biological and Human–Agent Motion: Functional Magnetic Resonance Imaging and Neuropsychological Evidence. The Journal of Neuroscience 33:15442–15453 Available at: http://dx.doi.org/10.1523/jneurosci.5868-12.2013.

Hardwick RM, Caspers S, Eickhoff SB, Swinnen SP (2018) Neural correlates of action: Comparing meta-analyses of imagery, observation, and execution. Neurosci Biobehav Rev 94:31–44.

Heider F, Simmel M (1944) An Experimental Study of Apparent Behavior. Am J Psychol 57:243–259.

Iacoboni M, Dapretto M (2006) The mirror neuron system and the consequences of its dysfunction. Nat Rev Neurosci 7:942–951.

Iacoboni M, Molnar-Szakacs I, Gallese V, Buccino G, Mazziotta JC, Rizzolatti G (2005) Grasping the intentions of others with one’s own mirror neuron system. PLoS Biol 3:e79.

Isik L, Koldewyn K, Beeler D, Kanwisher N (2017) Perceiving social interactions in the posterior superior temporal sulcus. Proc Natl Acad Sci U S A 114:E9145–E9152.

Karakose-Akbiyik S, Caramazza A, Wurm MF (2023) A shared neural code for the physics of actions and object events. Nat Commun 14:1–13.

Konkle T, Caramazza A (2013) Tripartite organization of the ventral stream by animacy and object size. J Neurosci 33:10235–10242.

Lacadie CM, Fulbright RK, Arora J, Constable RT, Papademetris X (2008) Brodmann Areas defined in MNI space using a new Tracing Tool in BioImage Suite. Human Brain Mapping, 2008. In. Human Brain Mapping.

Lee Masson H, Isik L (2021) Functional selectivity for social interaction perception in the human superior temporal sulcus during natural viewing. Neuroimage 245:118741.

Liu S, Ullman TD, Tenenbaum JB, Spelke ES (2017) Ten-month-old infants infer the value of goals from the costs of actions. Science 358:1038–1041.

McAleer P, Pollick FE, Love SA, Crabbe F, Zacks JM (2014) The role of kinematics in cortical regions for continuous human motion perception. Cogn Affect Behav Neurosci 14:307– 318.

Michotte A (1946) La perception de la causalité. (Etudes Psychol.), Vol. VI. 296 Available at: https://psycnet.apa.org/fulltext/1948-01004-000.pdf.

Mulliken GH, Musallam S, Andersen RA (2008) Decoding trajectories from posterior parietal cortex ensembles. J Neurosci 28:12913–12926.

Oosterhof NN, Connolly AC, Haxby JV (2016) CoSMoMVPA: Multi-Modal Multivariate Pattern Analysis of Neuroimaging Data in Matlab/GNU Octave. Front Neuroinform 10:27.

Opfer JE (2002) Identifying living and sentient kinds from dynamic information: the case of goal-directed versus aimless autonomous movement in conceptual change. Cognition 86:97–122.

Papademetris X, Jackowski MP, Rajeevan N, DiStasio M, Okuda H, Constable RT, Staib LH (2006) BioImage Suite: An integrated medical image analysis suite: An update. Insight J 2006:209.

Patri J-F, Cavallo A, Pullar K, Soriano M, Valente M, Koul A, Avenanti A, Panzeri S, Becchio C (2020) Transient disruption of the inferior parietal lobule impairs the ability to attribute intention to action. Curr Biol 30:4594–4605.e7.

Peelen MV, Downing PE (2017) Category selectivity in human visual cortex: Beyond visual object recognition. Neuropsychologia 105:177–183.

Pelphrey KA, Morris JP, McCarthy G (2004) Grasping the intentions of others: the perceived intentionality of an action influences activity in the superior temporal sulcus during social perception. J Cogn Neurosci 16:1706–1716.

Pitcher D, Ungerleider LG (2021) Evidence for a Third Visual Pathway Specialized for Social Perception. Trends Cogn Sci 25:100–110.

Pramod RT, Cohen MA, Tenenbaum JB, Kanwisher N (2022) Invariant representation of physical stability in the human brain. Elife 11 Available at: http://dx.doi.org/10.7554/eLife.71736.

R Core Team (2021) R: A language and environment for statistical computing. R Foundation for Statistical Computing, Vienna, Austria. Vienna, Austria.

Rizzolatti G, Craighero L (2004) The mirror-neuron system. Annu Rev Neurosci 27:169–192.

Saxe R, Xiao D-K, Kovacs G, Perrett DI, Kanwisher N (2004) A region of right posterior superior temporal sulcus responds to observed intentional actions. Neuropsychologia 42:1435–1446.

Scholl BJ, Gao T (2013) Perceiving Animacy and Intentionality. In: Social Perception, pp 197–230. The MIT Press.

Scholl BJ, Tremoulet PD (2000) Perceptual causality and animacy. Trends Cogn Sci 4:299–309.

Schultz J, Frith CD (2022) Animacy and the prediction of behaviour. Neurosci Biobehav Rev 140:104766.

Schwettmann S, Tenenbaum JB, Kanwisher N (2019) Invariant representations of mass in the human brain. Elife 8 Available at: http://dx.doi.org/10.7554/eLife.46619.

Sliwa J, Freiwald WA (2017) A dedicated network for social interaction processing in the primate brain. Science 356:745–749.

Tarhan L, Konkle T (2020) Sociality and interaction envelope organize visual action representations. Nat Commun 11:3002.

Tremoulet PD, Feldman J (2006) The influence of spatial context and the role of intentionality in the interpretation of animacy from motion. Percept Psychophys 68:1047–1058.

Urgesi C, Candidi M, Avenanti A (2014) Neuroanatomical substrates of action perception and understanding: an anatomic likelihood estimation meta-analysis of lesion-symptom mapping studies in brain injured patients. Front Hum Neurosci 8:344.

Van Nuenen BF, Kuhtz-Buschbeck J, Schulz C, Bloem BR, Siebner HR (2012) Weightspecific anticipatory coding of grip force in human dorsal premotor cortex. J Neurosci 32:5272– 5283.

Watson CE, Cardillo ER, Ianni GR, Chatterjee A (2013) Action concepts in the brain: an activation likelihood estimation meta-analysis. J Cogn Neurosci 25:1191–1205.

Wurm MF, Caramazza A (2022) Two “what” pathways for action and object recognition. Trends Cogn Sci 26:103–116.

Wurm MF, Caramazza A, Lingnau A (2017) Action Categories in Lateral Occipitotemporal Cortex Are Organized Along Sociality and Transitivity. J Neurosci 37:562–575.

Yildirim I, Wu J, Kanwisher N, Tenenbaum J (2019) An integrative computational architecture for object-driven cortex. Curr Opin Neurobiol 55:73–81.

Zacks JM, Swallow KM, Vettel JM, McAvoy MP (2006) Visual motion and the neural correlates of event perception. Brain Res 1076:150–162.

