## Supplementary Information for "The role of agentive and physical forces in the neural representation of motion events"

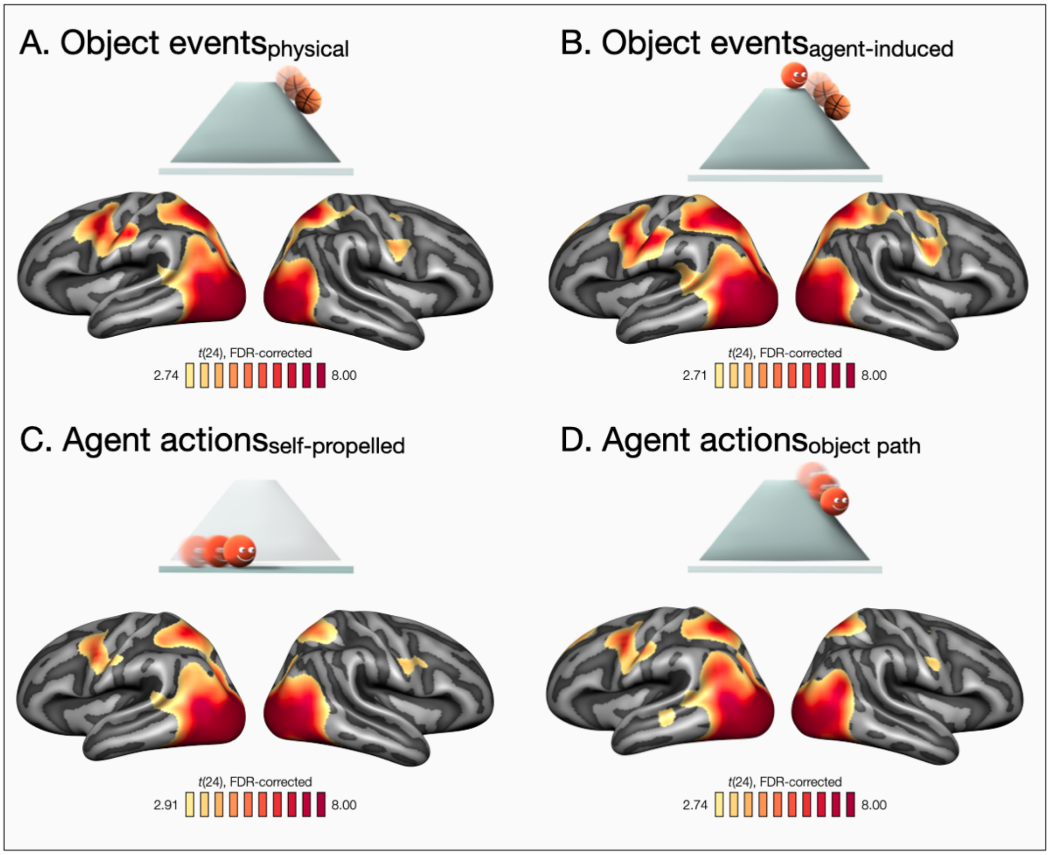


**Supplementary Figure 1 - Whole-brain maps depicting univariate responses to different experimental conditions against baseline.** (**A-D**) Compared against the baseline, all experimental conditions led to increased activity in multiple regions spanning occipital, posterior temporal, frontal, and parietal cortices. All maps are FDR-corrected.

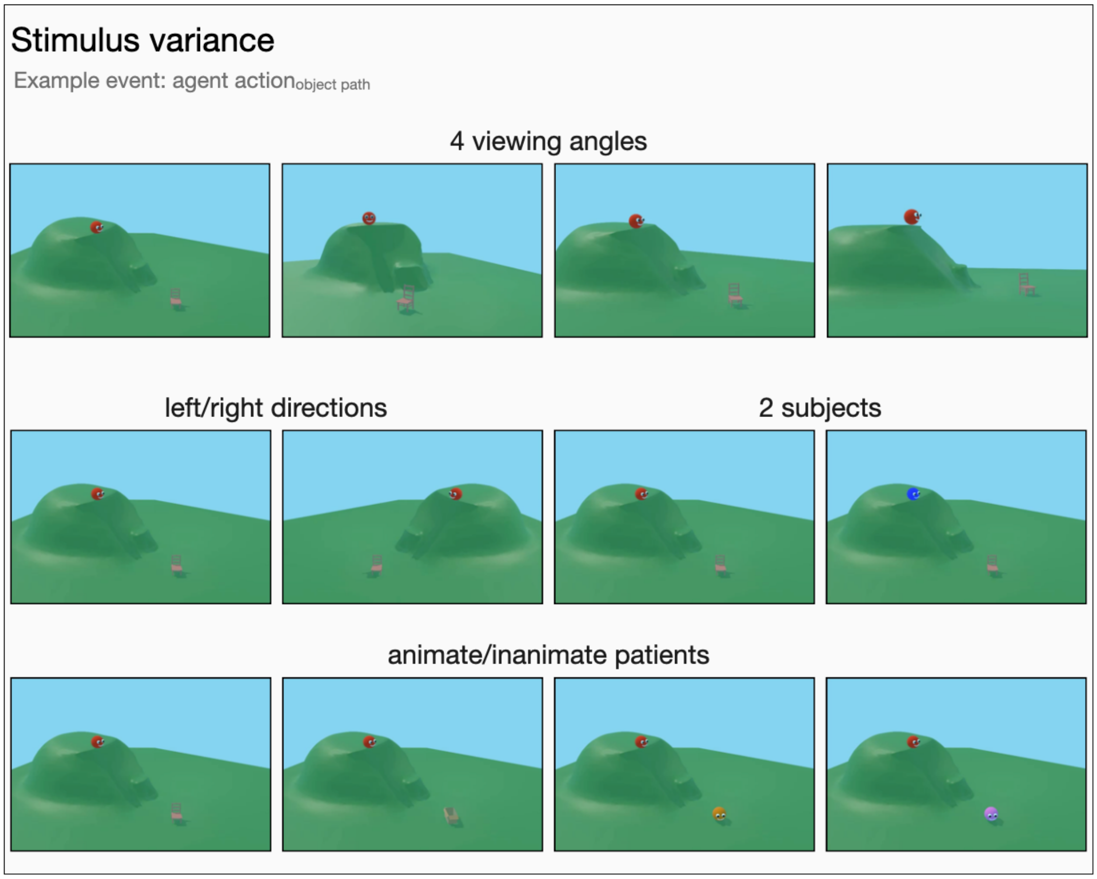


**Supplementary Figure 2 – Stimulus variance.** Still images depicting different exemplars that were generated for each motion trajectory and experimental condition by varying viewing angle, direction of movement, subjects, and passive patients. For this display, agent $action_{object\text{-}path}$ was used as an example. For instance, the 'hit' event for agent $action_{object\text{-}path}$ would be presented across 64 exemplars that depict the same event through four viewing angles, from left and right directions, blue or red agents, and animate or inanimate passive patients (e.g., chair, box, orange agent, purple agent).

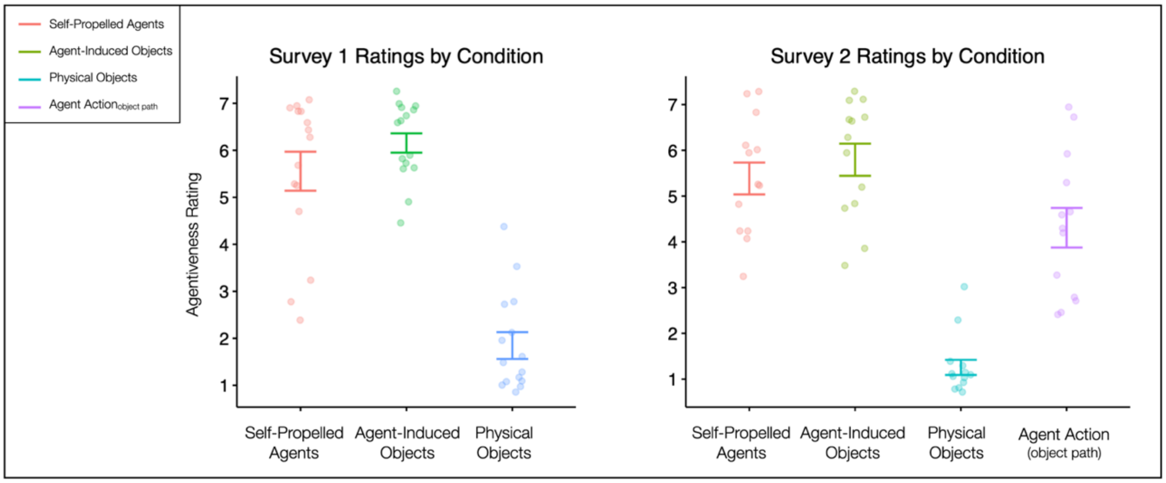


**Supplementary Figure 3 – Behavioral survey results.** An independent group of participants were asked to rate the degree to which the videos appeared agentive on a 7-point Likert scale (1 = not at all agentive, 7 = highly agentive). Two different surveys were used with or without the agent action$_{\text{object-path}}$ condition. Error bars indicate standard error of the mean (SEM), and individual participants are presented with colored dots.

### Supplementary Note 1 – Behavioral survey results

We collected a behavioral survey from an independent group of participants to validate certain aspects of our stimuli. For physical object events, we wanted to ensure that observers did not ascribe any agent involvement. On the other hand, for conditions that depicted agent movement (self-propelled agent actions, agent actions$_{\text{object-path,}}$ agent-induced object events), we wanted to ensure that observers endorsed the role of agents in shaping the event dynamics. This concern was less relevant for self-propelled agent actions since they depicted complete agentive control. However, for agent-induced object events, we wanted to ensure that the agent hitting the object is perceived as the causer of the object's motion. As for agent actions$_{\text{object-path}}$, the motion trajectories were driven by the physics engine and the agent depicted a sliding movement. Thus, there was a chance that observers would not interpret these events as sufficiently agentive. This concern was compounded by a potential spillover effect: if agent actions$_{\text{object path}}$ were seen as non-agentive, subjects might come to perceive the self-propelled agent actions as less agentive as well, given that they were carried out by identical "agents": spherical objects with eyes and a mouth.

To address these issues, we conducted two surveys in which respondents viewed videos from different conditions and were asked to rate the degree to which they appeared agentive, on a 7-point Likert scale (1 = not at all agentive; 7 = highly agentive). We clarified that "*Agentive events are events which could not have happened without the involvement of an animate agent. For instance, a soccer player kicking a goal is an agentive event, since the kick was initiated by an animate agent; in contrast, a leaf falling from a tree is not agentive, since the event was not initiated by an animate agent*." To test for a potential spillover effect, participants were randomly assigned to either Survey 1, which contained the self-propelled agent action conditions, physical object events, and agent-induced object events (n = 15); or Survey 2, which also contained the agent actions$_{\text{object-path}}$ condition (n = 13).

A repeated-measures ANOVA revealed a significant difference in agentiveness ratings across different experimental conditions (Survey 1: $F(2,28) = 50.72, p < 0.001, \eta_g 2 = 0.72$; Survey 2: $F(2, 25) = 56.72$, $p < 0.001$, $\eta_g 2 = 0.70$). Two-tailed pairwise t-tests showed that conditions that involved agents (i.e., agent $\text{actions}_{\text{self-propelled}}$, agent $\text{actions}_{\text{object-path}}$, object $\text{events}_{\text{agent-induced}}$) were perceived as significantly more agentive than physical object events (all *p*s < 0.001). Agentiveness ratings were high for all motion events that involved an agent: self-propelled agent actions (Survey 1: *Median* = 6.33, *SD* = 1.61; Survey 2: *Median* = 5.33, *SD* = 1.25), agent-induced object events (Survey 1: *Median* = 6.33, *SD* =.79; Survey 2: *Median* = 6.00, *SD* = 1.27), and agent $\text{actions}_{\text{object path}}$ (Survey 2: *Median* = 4.33, *SD* = 1.56). Participants did not ascribe agent involvement to physical object events (Agentiveness Rating - Survey 1: $Median_{physical\ objects} = 1.33$, $SD_{physical\ objects} = 1.10$; Survey 2: $Median_{physical\ objects} = 1.00$, $SD_{physical\ objects} = .60$).

An independent samples t-test between the self-propelled agent conditions in Survey 1 and Survey 2 did not show a significant difference in perception of agentive control ($t(26) = 0.31, p = .759, d = .12$), ruling out a spillover effect. In other words, self-propelled agent actions were perceived as similarly agentive regardless of whether they were accompanied by agent $\text{actions}_{\text{object path}}$ that depicted less agentive control. Note that in Survey 2 self-propelled agent actions were rated as more agentive than agent $\text{actions}_{\text{object-path}}$ ($t(12) = 4.07, p = .002, d = 1.13$), which is expected given that these stimuli presented an agent sliding down a hill as controlled by the physics engine. We chose to include both conditions in the experiment, given the lack of a spillover effect and the fact that the physical object events were always rated lower than events involving agents.

Overall, our survey results validated the use of our stimuli, ensuring that even relatively minor visual differences between spherical agents and objects nonetheless resulted in significantly different levels of perceived agency.

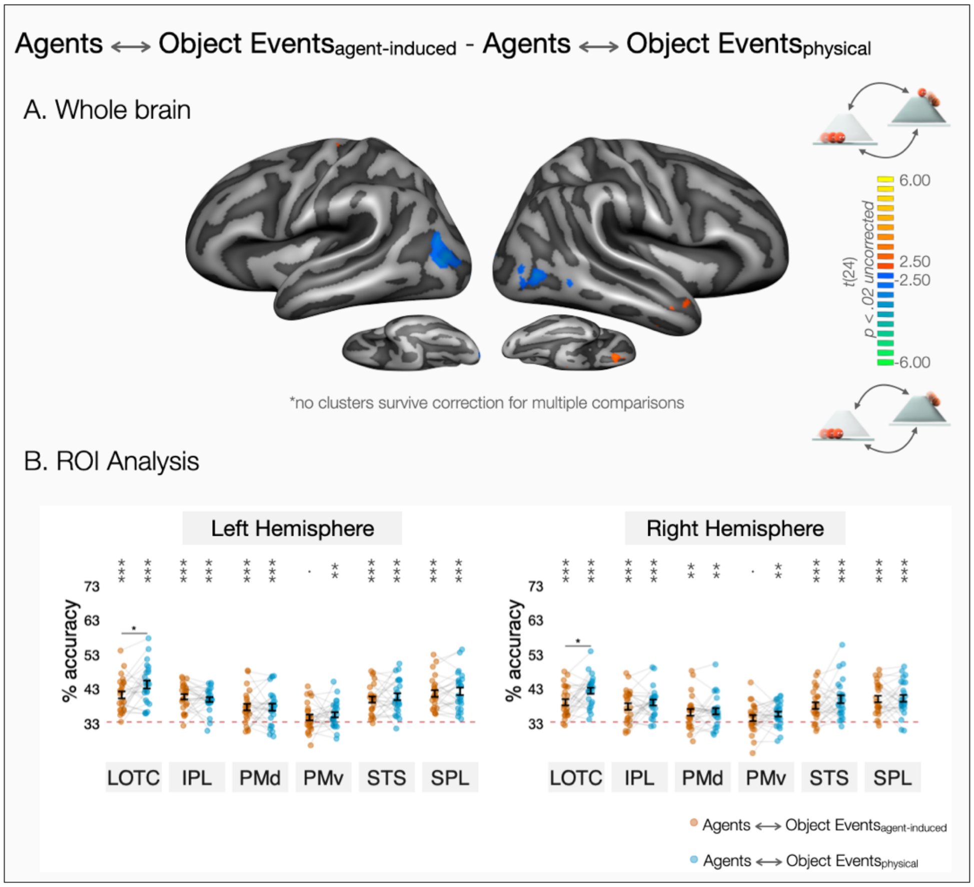


**Supplementary Figure 4 – Comparison of cross-animacy decoding as a function of causal agency (A)** Whole-brain contrast of cross-decoding between self-propelled agent actions and agent-induced object events or physical object events. No clusters survived correction for multiple comparisons. The map is thresholded at $p < .02$ to demonstrate significant differences that do not survive correction for multiple comparisons. (**B**) ROI decoding accuracies for cross-decoding between self-propelled agent actions and agent-induced object events or physical object events. Error bars indicate standard error of the mean (SEM), and asterisks indicate FDR-corrected effects of one-tailed t-tests for comparisons against chance-level (33.33%, $*p < .05$, $**p < 0.01$, $***p < 0.001$). Individual participants are connected via light gray lines. FDR-corrected pairwise two-tailed tests of estimated marginal means revealed stronger cross-decoding between self-propelled agent actions and physical object events in left and right LOTC ($*p < .05$, $**p < 0.01$, $***p < 0.001$).

**Supplementary Note 2 – Cross-animacy decoding ROI results and the role of causal agency.**

Various frontoparietal and posterior temporal brain regions showed successful cross-decoding between self-propelled agent actions and physical or agent-induced object events (see Figure 2A-B). Self-propelled agent actions and agent-induced object events are both tied to an agent cause, while physical object events are not. Does shared causal agency between self-propelled agent actions and agent-induced object events lead to better cross-animacy generalization in the neural representation of event dynamics? To test this question, we compared the cross-decoding strengths self-propelled agent actions and agent-induced object events with that of self-propelled agent actions and physical object events. In the whole brain, a two-tailed t-test did not lead to reliable differences in cross-decoding strength (Supplementary Figure 4A).

To obtain a finer grained understanding of differences in cross-decoding across specific regions, we extracted classification accuracies from independently defined regions of interest (ROIs) linked to action observation: lateral occipitotemporal cortex (LOTC), inferior parietal lobule (IPL), dorsal premotor cortex (PMd), ventral premotor cortex (PMv), posterior superior temporal sulcus (pSTS), and superior parietal lobule (SPL, see Methods for more details on ROI selection). All ROIs in both left and right hemispheres showed above chance cross-decoding of the three motion events across self-propelled agent actions and physical object events (see Supplementary Figure 4B). As for cross-decoding between self-propelled agent actions and agent-induced object events, all ROIs showed above chance cross-decoding except for ventral premotor cortex, which bordered at significance (see Supplementary Figure 4B, $p_{\text{left PMv}} = .06$, $p_{\text{right PMv}} = .09$).

In the ROI-analysis, we observed an ROI by event type interaction both in the left ($\chi2[5] = 18.67$, $p = .002$, $\Delta AIC = 8.66$) and the right hemispheres ($\chi2[5] = 17.62$, $p = .003$, $\Delta AIC = 7.62$). Cross-decoding between self-propelled agent actions and physical object events was stronger than that between self-propelled agent actions and agent-induced object events in left LOTC ($b = -3.02$, $p = .044$, $d = -.51$) and right LOTC ($b = -3.37$, $p = .013$, $d = -.62$). No significant difference was observed in other ROIs. At this juncture, we would like to note that agent-induced object events had a more complex event structure and depicted instrumental actions, whereas physical object events and self-propelled agent actions did not. This difference in complexity and event structure might explain stronger cross-decoding of self-propelled agent actions to physical object events compared to that between self-propelled agent actions and agent-induced object events and might have masked any possible contribution of causal agency to cross-animacy generalization. Further research is needed to fully understand the contributions of causal agency on the neural representation of event dynamics.

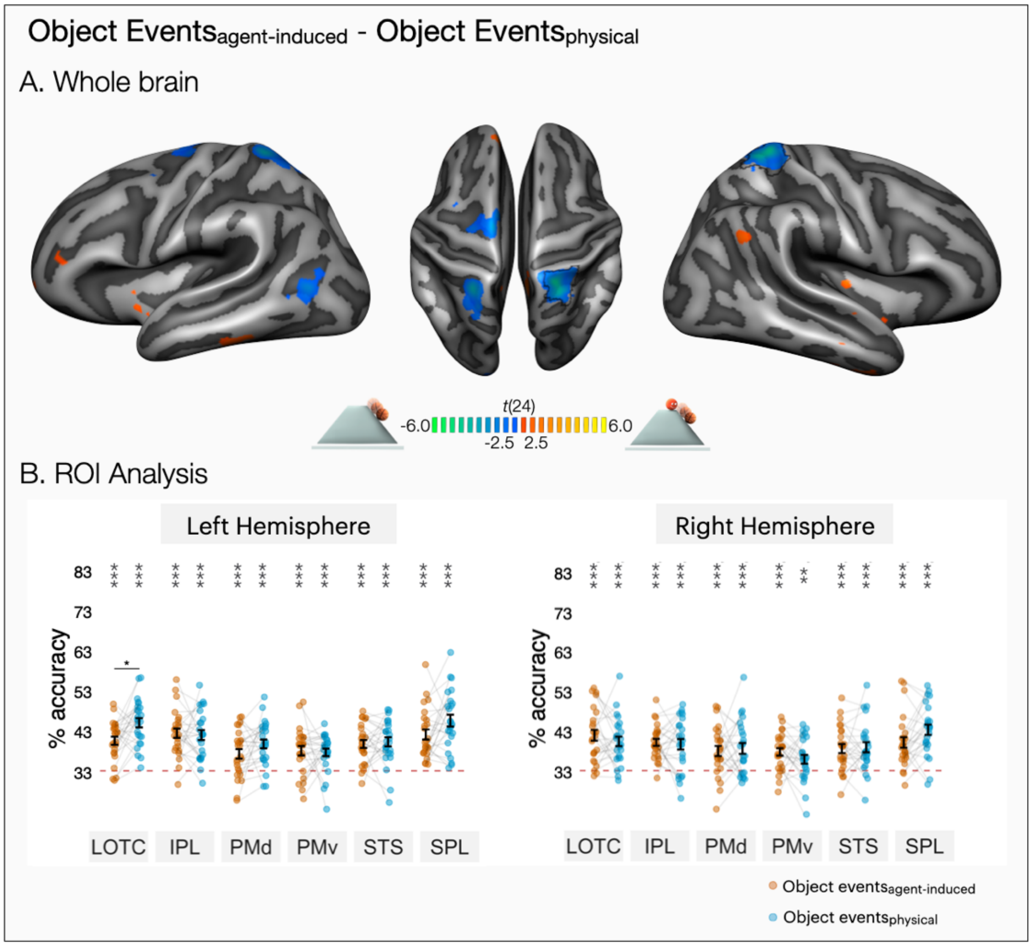


**Supplementary Figure 5 – Decoding contrast of agent-induced and physical object events. (A)** Whole-brain decoding contrast of agent-induced object events and physical object events. The map is thresholded at $p < .02$ to demonstrate significant differences that do not survive correction for multiple comparisons. Regions that survive correction for multiple comparison are highlighted with a black boundary (see the cluster in right superior parietal lobule). **(B)** ROI decoding accuracies for agent-induced object events and physical object events. Error bars indicate standard error of the mean (SEM), and asterisks indicate FDR-corrected effects of one-tailed t-tests for comparisons against chance-level (33.33%, *$p < .05$, **$p < 0.01$, ***$p < 0.001$). Individual participants are connected via light gray lines. FDR-corrected pairwise two-tailed tests of estimated marginal means showed better decoding of physical object events in left LOTC (*$p < .05$, **$p < 0.01$, ***$p < 0.001$).

**Supplementary Note 3 - Decoding contrast of agent-induced and physical object events.**

As an additional exploratory analysis to address the contributions of causal agency to the neural representation of event dynamics, we compared the decoding strengths of agent-induced object events with physical object events. In theory, regions that show stronger decoding of agent-induced object events compared to physical object events might be sensitive to causal agency behind an object's movement. In the whole-brain, physical object events were decoded at a higher accuracy than agent-induced object events in a cluster in right superior parietal lobule (Supplementary

Figure 5A). Additional clusters in left dorsal premotor cortex and superior parietal lobule showed a similar effect (*p*s < .005), but these clusters did not survive correction for multiple comparisons in the whole brain.

In the ROI analysis, we observed an ROI by event type interaction both in the left ($\chi2[5] = 24.80$, $p < .001$, $\Delta AIC = 14.80$) and right hemispheres ($\chi2[5] = 22.64$, $p < .001$, $\Delta AIC = 12.64$). Post-hoc contrasts revealed stronger decoding of physical object events compared to agent-induced object events in left LOTC ($b = -4.49$, $p = .017$, $d = -.64$), and a marginal trend in left SPL ($b = -3.46$, $p = .064$, $d = -.41$). The rest of the left hemisphere ROIs did not show a significant difference between agent-induced object events and physical object events. In the right hemisphere, decoding of physical and agent-induced object events were comparable in all ROIs except for right SPL where a medium sized effect did not survive correction for multiple comparisons ($b = -3.19$, $p = .173$, $d = -.45$).

Overall, in comparing object events with or without causal agency, we found right SPL to be more sensitive to events that were shaped by the inherent physics of the scene. However, note that agent-induced object events involved a more complex event structure, in which three entities were involved: in addition to an agent and a passive patient, an instrument (i.e., the ball being pushed by the agent). Furthermore, with the involvement of an agent, people might have processed these scenes differently, also paying attention to the agent throughout the videos. As such, this condition raises a variety of issues that could impact differences in neural activity patterns and mask the potential effects of causal agency, which are beyond the scope of this study.
